# Rational Design and Modeling of Auxetic Fiber Scaffolds for Soft Tissue Engineering via Melt Electrowriting

**DOI:** 10.1101/2025.05.15.654200

**Authors:** Óscar Lecina-Tejero, Jirawat Iamsamang, Pilar Alamán-Díez, Elena García-Gareta, Jesús Cuartero, María Ángeles Pérez, Miguel Castilho, Carlos Borau

**Author notes:** These authors contributed equally to this work (Óscar Lecina-Tejero, Jirawat Iamsamang).

## Abstract

**Background:** The development of regenerative biomaterial scaffolds with mechanical properties tailored to native soft tissues remains a challenge in tissue engineering. Mechanical metamaterials with auxetic architectures have taken enormous flight over the past years, as they offer improved control over mechanical function and performance through spatially defined material architecture. However, despite their potential, auxetic scaffold designs are often developed empirically, and the relationship between scaffold microarchitecture, mechanical function, and manufacturability remains poorly defined, limiting their applicability in tissue engineering. In this study, we present a computational framework for the rational design of auxetic fiber scaffolds manufactured using Melt Electro-writing (MEW) for soft tissue engineering. To address the challenge of tailoring scaffold mechanical properties to native tissue, we integrated finite element method (FEM) modeling with experimental validation to predict the mechanical behavior of the scaffolds.

**Method:** The micro-fibrous scaffolds with customizable auxetic re-entrant structures were designed using Python scripts for generating G-code used in fabrication and for model definition in FEM modeling, respectively. Four distinct auxetic structures – HCELL, SREG, SINV, and STRI – were selected for this study to cover the wide range of mechanical properties of soft tissue, and their geometry parameters were adjusted to reduce the pore size while considering printing resolution. FEM modeling was employed to predict the mechanical behavior of the scaffolds. To validate the predictions, the scaffolds were manufactured via MEW and characterized mechanically under biaxial tensile loading.

**Results and conclusion:** MEW manufacturing enabled the production of auxetic fiber scaffolds with sub-millimeter pore resolution, while rational design allowed to correct fabrication discrepancies and improve printed design fidelity. Experimental biaxial characterization of the scaffolds revealed design-dependent, biphasic, non-linear mechanical response, and the developed FEM model was capable of not only accurately simulating the observed mechanical response but also accounting for variations introduced by MEW during fabrication. Additionally, auxetic scaffolds supported fibroblast viability over 7 days, confirming biocompatibility as a first step toward soft tissue engineering. The good alignment between numerical predictions and experimental results highlights the value of FEM modeling as a powerful computational tool for guiding the design and optimization of auxetic scaffolds with tailored mechanical performance for specific soft tissue requirements. The presented methodology offers a significant advancement in the development of functional biomaterials for biomedical applications.

## 1. Introduction

Soft tissues in the human body exhibit a wide range of mechanical properties that play essential roles in structural support, protection, and function. When damaged by trauma, disease, or surgery, their ability to regenerate may be limited, necessitating the use of engineered scaffolds to support and guide tissue repair[1]. Traditional treatments, such as autologous grafts, often face limitations including tissue availability, donor compatibility issues, and tissue morbidity responses[2–4]. To overcome these challenges, recent advances in tissue engineering have explored the use of micro-structured scaffolds fabricated through technologies like electrospinning or melt electro-writing (MEW). These approaches aim to recreate the structural and mechanical characteristics of the extracellular matrix (ECM), offering improved conditions for tissue regeneration[5].

A critical factor in soft tissue repair is the local mechanical environment. Mechanical signals influence cellular behaviors such as proliferation, migration, and differentiation, all of which are necessary for effective healing. Excessive or misdirected mechanical stimuli can disrupt angiogenesis, promote fibrosis, or alter the phenotype of cells like fibroblasts, potentially leading to suboptimal tissue regeneration[6]. Therefore, scaffolds must not only provide structural support but also exhibit mechanical properties that are compatible with the surrounding tissue environment.

One of the main challenges in developing such scaffolds lies in the broad mechanical diversity of soft tissues, from the elasticity of dermal skin layers to the more fibrous nature of tendons or fascia. Mimicking the fibrous architecture and biomechanical response of the native ECM – particularly the role of collagen and elastin – has led to the emergence of fiber-based scaffolds. These constructs aim to replicate microscale ECM features and allow for tuning of stiffness, elasticity, and anisotropy by manipulating fiber orientation, spacing, and geometry[7]. A growing body of research has explored micro-fibrous scaffold design for soft tissue applications[5,8–11], including the integration of auxetic structures, which offer promising mechanical characteristics for dynamic environments[12–17].

Auxetic metamaterials exhibit a negative Poisson’s ratio, expanding laterally when stretched and contracting when compressed. This counterintuitive behavior enables enhanced flexibility, greater energy absorption, and uniform stress distribution across complex deformation states[18,19]. Such properties are advantageous in applications requiring compliance and adaptability, including the repair of tissues subjected to multidirectional strain[20]. Auxetic mechanics have been observed in natural tissues[21–27] and are governed not only by material composition but also by internal geometry[28,29]. Fiber-based scaffolds offer a versatile platform for implementing auxetic designs such as re-entrant and chiral patterns[30–32], allowing control over mechanical performance including strain distribution and deformation modes, which makes them suitable for tailoring scaffold behavior to match specific tissue requirements[33,34].

Among the available fabrication techniques, MEW offers unique advantages for developing scaffolds that closely mimic native ECM architecture. This solvent-free process enables the precise deposition of micro-to submicron-scale fibers utilizing electrically driven extrusion of molten polymers. The result is highly organized scaffolds with programmable architectures and fine-tuned mechanical properties. MEW can produce both flat and tubular constructs with defined porosity, anisotropy, and mechanical responses, including auxetic behavior[35–40]. Polycaprolactone (PCL), a biodegradable and biocompatible polymer, is well suited for MEW due to its processability and prior use in various soft tissue engineering applications[41–43].

Despite these advancements, a systematic framework that efficiently combines numerical prediction, fabrication, and experimental characterization, linking scaffold design parameters to mechanical performance is necessary to improve the functionality by tailoring the scaffold design to specific mechanical properties of target soft tissues[20,33].

In this study, we present the development of auxetic PCL micro-fibrous scaffolds for soft tissue engineering, fabricated via MEW and informed by rational design principles. Our primary aim is to create computational tools capable of predicting the mechanical response of various scaffold geometries through finite element method (FEM) modeling, validated against experimental biaxial tensile testing. We further examine how deviations introduced during the printing process influence mechanical behavior and demonstrate that our FEM models can capture these effects. Skin tissue was selected as a reference due to its mechanical similarity to the auxetic scaffolds. Consequently, basic fibroblast culture assays were performed to evaluate the biological viability of these scaffolds (Supp. Info). However, the broader goal of this study is to establish a robust framework for designing MEW auxetic scaffolds that can be tailored for a wide range of soft tissue applications.

## 2. Materials and Methods

This section describes the approach employed in the design, fabrication, and evaluation of auxetic scaffolds. As illustrated in Figure 1, the methodology comprises five interconnected stages: auxetic design (section 2.1), computational modeling (section 2.2), scaffold fabrication (section 2.3), mechanical characterization (section 2.4), and cellular viability assessment (Supp. Info). The following subsections provide details of the specific analytical techniques, materials, and methods used at each stage.

**Figure 1.**
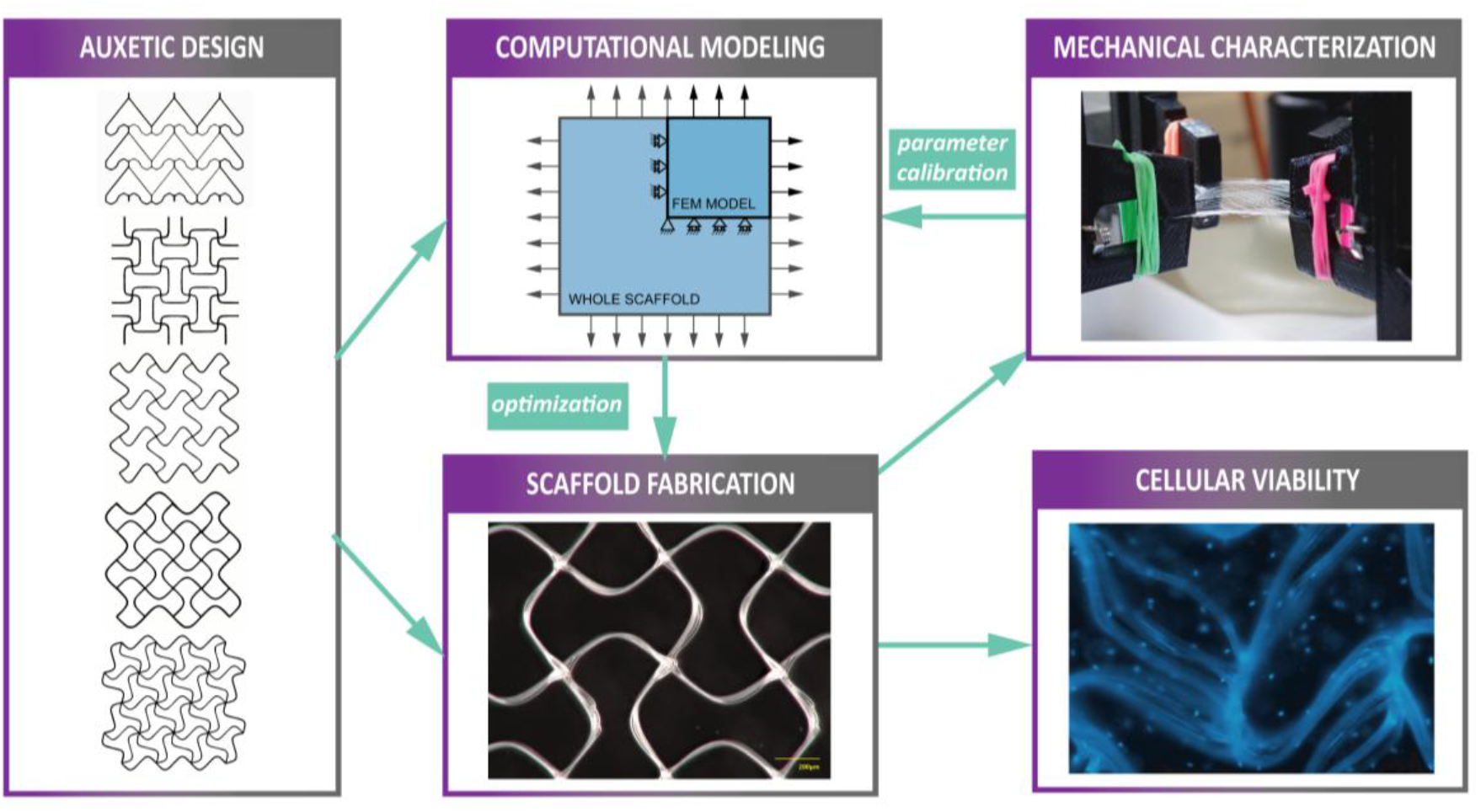
Schematic representation of the workflow, illustrating the sequential approach from auxetic design and computational modeling to auxetic scaffold melt electrowriting, mechanical characterization, and cellular viability assessment.

### 2.1 Design of auxetic scaffolds

Auxetic scaffold designs were conceived as 2D geometries based on interconnected fibers, stacked in identical layers to form 3D structures. Five re-entrant designs were selected, which are known to exhibit auxetic behavior[44–47]. The designs are designated as *Arrowhead, H-cell (HCELL), S-regular (SREG), S-inverted (SINV)*, and *S-triangular (STRI)*, as shown in Figure 2.

**Figure 2.**
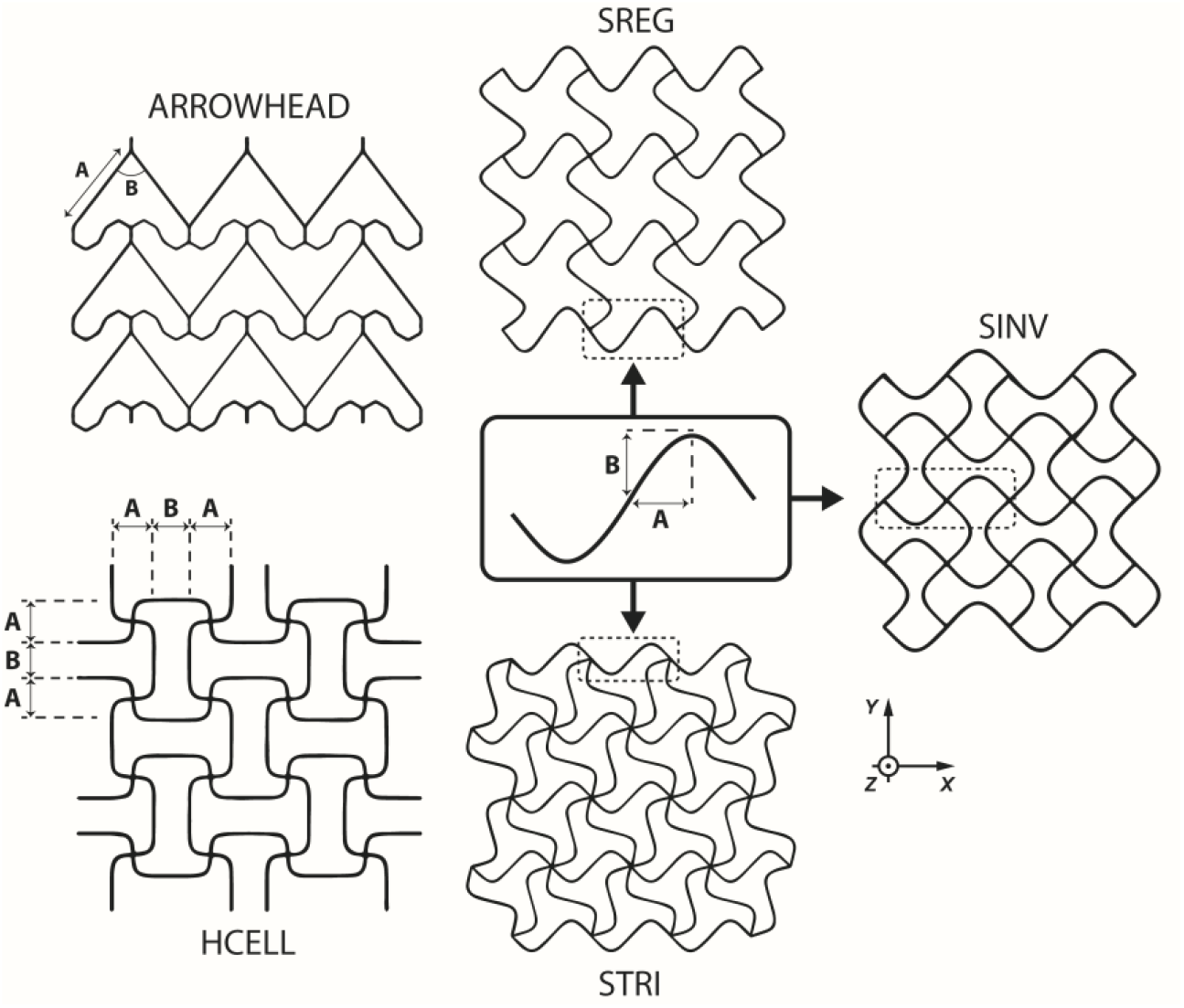
Scheme of the selected auxetic designs and their inner parameters.

A basic unit was defined for each auxetic design, which was then replicated in both the planar (x-y) and out-of-plane (z) dimensions to construct the scaffolds. Each basic unit was characterized by specific inner parameters, typically including fiber lengths in different directions. Additional assembly parameters were defined for the scaffold construction, comprising the number of basic unit replications in the x and y directions and the number of layers stacked in the z dimension to form the 3D scaffold. Table 1 summarizes the geometrical parameters selected for the numerical modeling of each auxetic design.

**Table 1:** Geometry parameters set for the numerical models reproducing the features of the printed scaffolds.

| Auxetic design | Parameter 'A' [$\mu\text{m}$] | Parameter 'B' [$\mu\text{m}$] | Fiber diameter [$\mu\text{m}$] | Width (X) [mm] | Height (Y) [mm] | Layers (Z) [-] |
| --- | --- | --- | --- | --- | --- | --- |
| *Arrowhead* | 1000 | 60 [°] | 12 | 5 | 5 | 10 |
| *HCELL* | 150 | 250 | 12 | 5 | 5 | 10 |
| *SREG* | 250 | 250 | 12 | 5 | 5 | 10 |
| *SINV* | 250 | 187.5 | 12 | 5 | 5 | 10 |
| *STRI* | 250 | 187.5 | 12 | 5 | 5 | 10 |

### 2.2 Computational modeling Numerical model and simulations

Numerical simulations were conducted using the finite element method (FEM) with ABAQUS software v6.14 (3DS Dassault Systems). Python-based ABAQUS scripting was utilized to automate model definition and result post-processing, enabling the creation and analysis of multiple models with varying design parameters (Table 1).

The fibers were modeled using 3D beam linear elements (B31), with mesh controls over the length-to-diameter ratio of the elements, setting a value greater than 10 for all the elements to avoid numerical analysis distortions caused by the definition of the beam element dimensions. The beam element simplification is valid since the fibers are expected to experience stretching, eliminating concerns of buckling. Fiber layers were interconnected using node coupling in the nodes near the fiber joint zones to create a simplified 3D model of the scaffolds that could reproduce the solidary behavior of the fibers stacked in layers. The material behavior of PCL was based on uniaxial tensile test data of bulk PCL (Figure S2 – Supp. Info). Thus, it was modeled with linear elastic behavior (E = 100 MPa) up to a yield strength of 12 MPa, followed by a progressively increasing stiffness in the plastic region.

FEM simulations of the biaxial tests were conducted using a quarter-symmetry model of the scaffolds (Figure 3), reducing the number of elements required for the numerical models of the scaffolds to a value of around 10000 elements. The numerical analysis was performed using Abaqus explicit solver with dynamic load steps, and symmetric boundary conditions were applied along the cut planes to replicate the scaffold’s behavior accurately. Prescribed displacements were imposed on nodes along the loading edges, while nodes on the constrained edges were fixed. Nodal displacements from the deformed edges and reaction forces from the constrained edges were extracted from simulations. Post-processing of the data allowed for the calculation of the mechanical properties, including effective stress and strain. For these calculations, the scaffold was treated as a homogeneous continuum, with its entire volume considered for length and cross-sectional area determinations.

**Figure 3.**
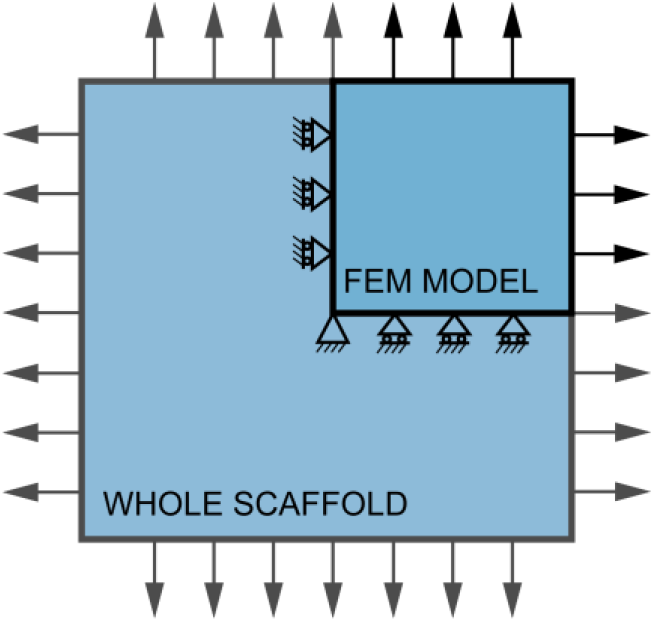
Boundary conditions prescribed for the numerical models of the scaffolds, replicating in a quarter-sized model the mechanical requirements of the experimental biaxial tensile tests performed with the printed scaffolds.

#### Python scripts

All computational processes in this study were executed in Python (Python 3.9). These scripts facilitated the key aspects of the research: (1) generation of G-code files for scaffold printing and (2) creation of the FEM models (ABAQUS inputs). The G-code generation script translates the auxetic designs into printer-readable instructions, ensuring precise fabrication of the scaffolds. The FEM scripts transform the physical dimensions and material properties into a numerical model including the necessary boundary conditions. These tools collectively enable a streamlined, reproducible workflow from design conception to data analysis. Further details are provided in Supp. Info. All the codes are publicly available on GitHub[48].

### 2.3 Scaffold fabrication

The PCL scaffolds were fabricated by the in-house MEW setup (Eindhoven University of Technology, Netherlands). This setup comprised a motion system (ToolChanger, E3D, UK) and a custom-made printhead whose linear motion was controlled via G-code language (Duet3D, RRF3) (Figure S1 – Supp. Info). The printing path of each auxetic design was constructed by the G-code sequences, generated by Python scripts. Before printing, medical-grade PCL (PC12, Corbion, Netherlands) was stored at 4°C and pre-warmed at 37°C before being loaded into a glass syringe (Fortuna® Optima®, Poulten & Graf, 1mL Luer lock) and preheated with reservoir temperatures of 100°C for at least an hour until no bubbles were observed. During printing process, preheated PCL was extruded through a 25G stainless-steel needle (Fisnar) by compressed air at approximately 1.2bar (Festo VPPE-2-1-1/8-6-010-E1). A high voltage between +3.5 and +4.0kV (Heinzinger, LNC 10000-5P) was applied at the stainless-steel needle. A printing speed of 200 mm/min was used. Prior to printing, key process parameters (dispensing pressure, melting temperature, high voltage, and collector speed) were monitored using a digital camera (AM4113ZTL, Dino-lite, Netherlands) to ensure stable deposition of fibers, minimizing coiling and meandering. The pore area, fiber diameter, and scaffold thickness were measured using a digital microscope (Keyence VHX-970FN). The scaffold’s pore size was defined by the minimal Ferret diameter of pores. The theoretical pore size and pore area were derived from the area of the pore from G-code positions subtracted by the area occupied by the surrounding fibers (Figure S2 – Supp. Info). The printing accuracy was determined by the percentage of measured values to the theoretical values of pore size and pore area.

### 2.4 Mechanical characterization

Scaffold mechanical characterization was performed with a biaxial tensile machine (CellScale® Biotester 5000). The scaffolds (12x12 mm^2^) were clamped with four holders in the XY plane (Figure 5-A). Scaffold thickness was measured with a digital microscope (Keyence VHX-970FN), taking at least 5 measurements per scaffold prior to testing. The tensile test started from a displacement of 0.1 to 36mm (edge-to-edge) at a strain rate of 0.1 mm/s. Tensile stress (σ) was calculated using Eq. 1, where force (F) was recorded from 2.5N load cells, thickness (T) was measured by the digital microscope, and the width (*W*) of the specimen was assumed to be the width of a tensile clamp (5 mm). Tensile strain was calculated using Eq. 2, where L_0_ is the initial displacement before testing and *L* is the displacement from the machine. The yield point was defined by a maximum ε in the linear part of the elastic region before plastic deformation identified by least-square regression (R^2^ > 0.97). The stiffness (*E*) was then calculated based on regression coefficients from this linear part, and strain energy density was derived from the area under the σ − *ϵ* curve to the yield point.

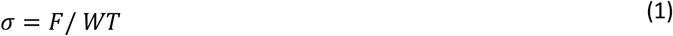

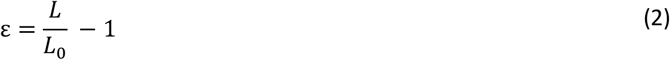

It is worth mentioning that *Arrowhead* designed scaffolds, initially subjected to uniaxial tensile tests (Figure S3 – Supp. Info), were excluded from further mechanical characterization as they could not withstand the biaxial conditions.

### Statistical analysis

For mechanical properties, statistical analysis was done by MATLAB 2023b (MathWorks™). All data were reported as average ± standard deviation. The normality and equal variance were examined using Shapiro-Wilk’s test and Bartlett’s test, respectively. For data with normal distribution, the differences between groups were detected by either one-way Brown–Forsythe’s ANOVA with Tukey’s post hoc (equal variance) or Welch’s ANOVA (unequal variance). Otherwise, Kruskal-Wallis tests were performed. The statistical significance level is 5% for all analyses (α=0.05).

## 3. Results and Discussion

### MEW fabrication of auxetic scaffolds requires optimization to achieve design-accurate geometries

Scaffolds with four distinct auxetic designs: *HCELL, SREG, SINV*, and *STRI* (Figure 4-A), were successfully fabricated with the design parameters described in Table 1. Due to the challenges in the MEW process for fabricating the scaffolds with small pores (<300 µm), e.g., fiber bridging[49] and residual electrostatic charges[50], these parameters were carefully selected and optimized to ensure that scaffolds were fabricated with fidelity while minimizing pore sizes for cell seeding and attachment, resulting in three different groups of pore sizes: small (*HCELL* and *SINV*), medium (*STRI*), and large (*SREG*). The fiber diameters of all scaffolds were maintained within the range of 10 - 17 µm. The statistical analysis revealed no significant differences in fiber diameter among the four designs (Figure 4-B). Moreover, the pore size (Figure 4-C) and pore area (Figure 4-D) of these scaffolds did not deviate significantly from the theoretical values (92 ± 3%, and 90 ± 4%, respectively). The pore area of *HCELL* (0.24 ± 0.01 mm^2^) was comparable to those of *SINV* scaffolds (0.20 ± 0.01 mm^2^). However, they are approximately 1.5 and 3.6 times smaller than those of *STRI* and *SREG*, respectively. The pore size showed a similar trend corresponding to the pore area. Likewise, the scaffolds with *HCELL* and *SINV* designs had similar pore sizes (0.46 ± 0.03 mm), which were 1.7 and 2.5 times smaller than those of the *STRI* and *SREG* designs, respectively.

**Figure 4.**
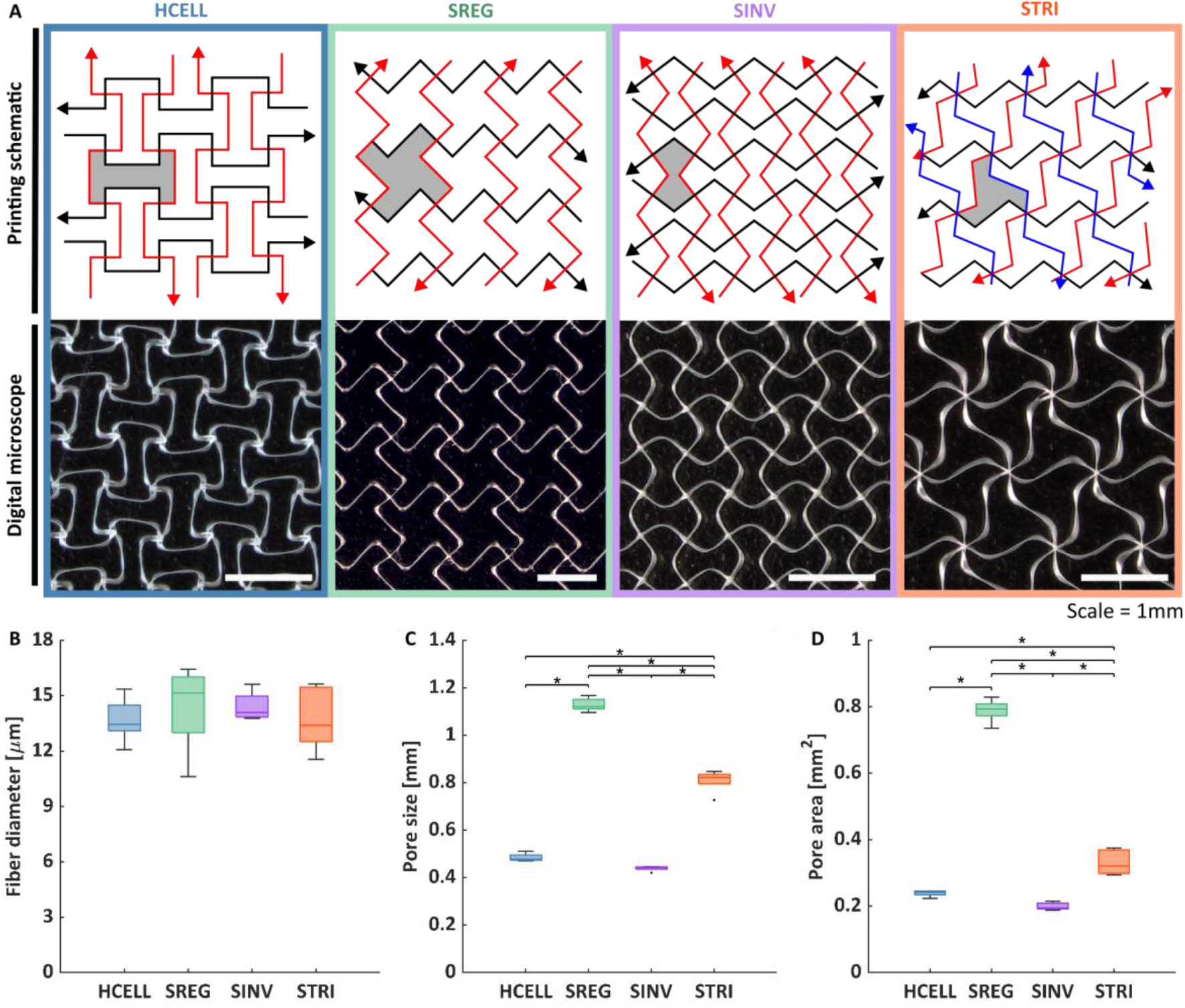
Printing schematic and fabricated scaffolds (A). The printing paths for each design are represented by arrows with black, red, and blue where the shaded areas (gray color) represent areas for quantifying pore area (B) and pore size (C) from the digital microscope images. The fiber diameters of fabricated scaffolds (D) have no statistical difference between designs, while the pore area and pore size show a significant difference (*p < 0.01), except those of SINV and HCELL (not significant: ns, p > 0.05).

**Figure 5.**
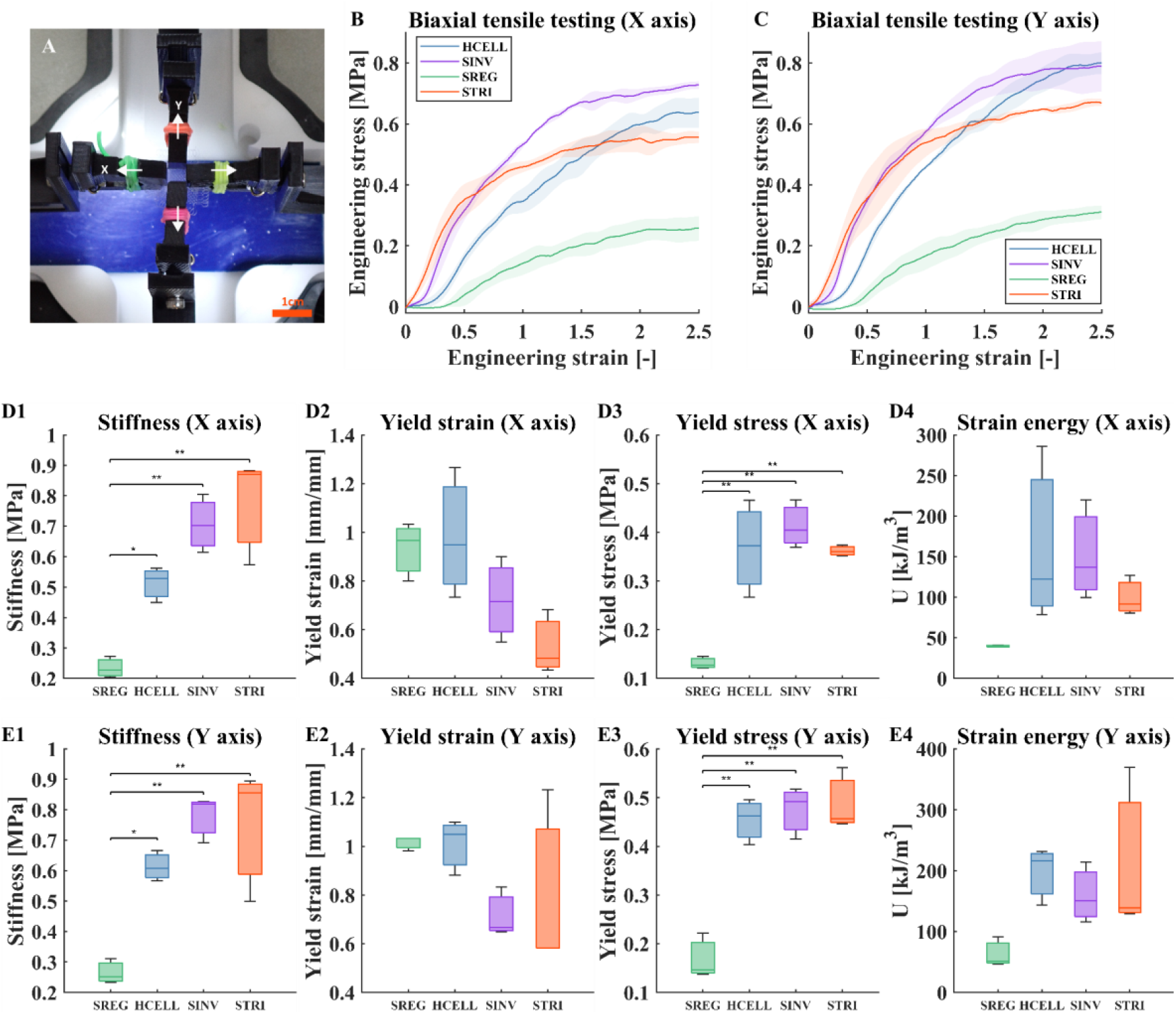
Mechanical properties of scaffolds with HCELL, SINV, SREG, and STRI designs. The biaxial tensile testing, where the white arrows represent directions of tensile testing (A) (scale bar = 1cm). The stress and strain curve in the X direction (B) and Y direction (C). Stiffness in the X and Y directions (D1, E1). Yield strain from least-squares regression in the X and Y directions (D2, E2) and corresponding yield stresses (D3, E3). Strain energy density up to the yield point in the X and Y directions (D4, E4). (*p < 0.05, **p < 0.01)

### Mechanical behavior of the scaffolds is strongly dependent on their auxetic fiber design

Once fabricated, the scaffolds were subjected to biaxial tensile tests, where they were clamped with four grips in the XY plane (Figure 5-A). No slipping of the specimen from the grips was observed during the tensile testing. The σ − *ϵ* curves in the X and Y axes showed similar trends in all four designs (Figure 5-B, C), with all the designs presenting a non-linear behavior consisting of an initial low-stress response, followed by a linear rise, and finally transitioning to a plateau at large strains, where plasticity, or even rupture, appeared within the scaffold fibers.

The yield stress and strain, however, were different between the auxetic designs. For example, the stiffness (Figure 5-D1, E1) of *SREG* in both X and Y axes (X axis: 0.23 ± 0.04 MPa; Y axis: 0.27 ± 0.04 MPa) was approximately 2 times lower than those of *HCELL* and 3 times lower than those of *SINV* and *STRI*. The yield stresses (Figure 5-D3, E3) of *SREG* in both axes (0.13 ± 0.01 MPa, 0.17 ± 0.04 MPa) were also the lowest among those of the other designs, being around 2.6 – 3.2 times lower than those of *HCELL, SINV*, and *STRI*. However, due to unequal variances and the limited number of replicates, no significant statistical differences in yield strain (Figure 5-D2, E2) and strain energy density (Figure 5-D4, E4) were observed among the four designs (Chi-square(3.8)=4.0-7.6, p=0.06-0.27), even though these differences were visually noticeable (Figure 5-D2, E2, and Figure 5-D4, E4).

Images collected from the biaxial tensile machine allowed us to observe the auxetic behavior of the scaffolds’ structure over the increasing values of strain (Figure 6). The basic cells of the scaffold gradually unfolded from the initially auxetic designs up to a point where they reached full extension and no longer exhibited auxetic mechanics. This point varied slightly between experiments and designs but occurred consistently around strain values between 0.4 and 0.6. Henceforth, we will refer to this point as the functional limit of the scaffolds.

**Figure 6.**
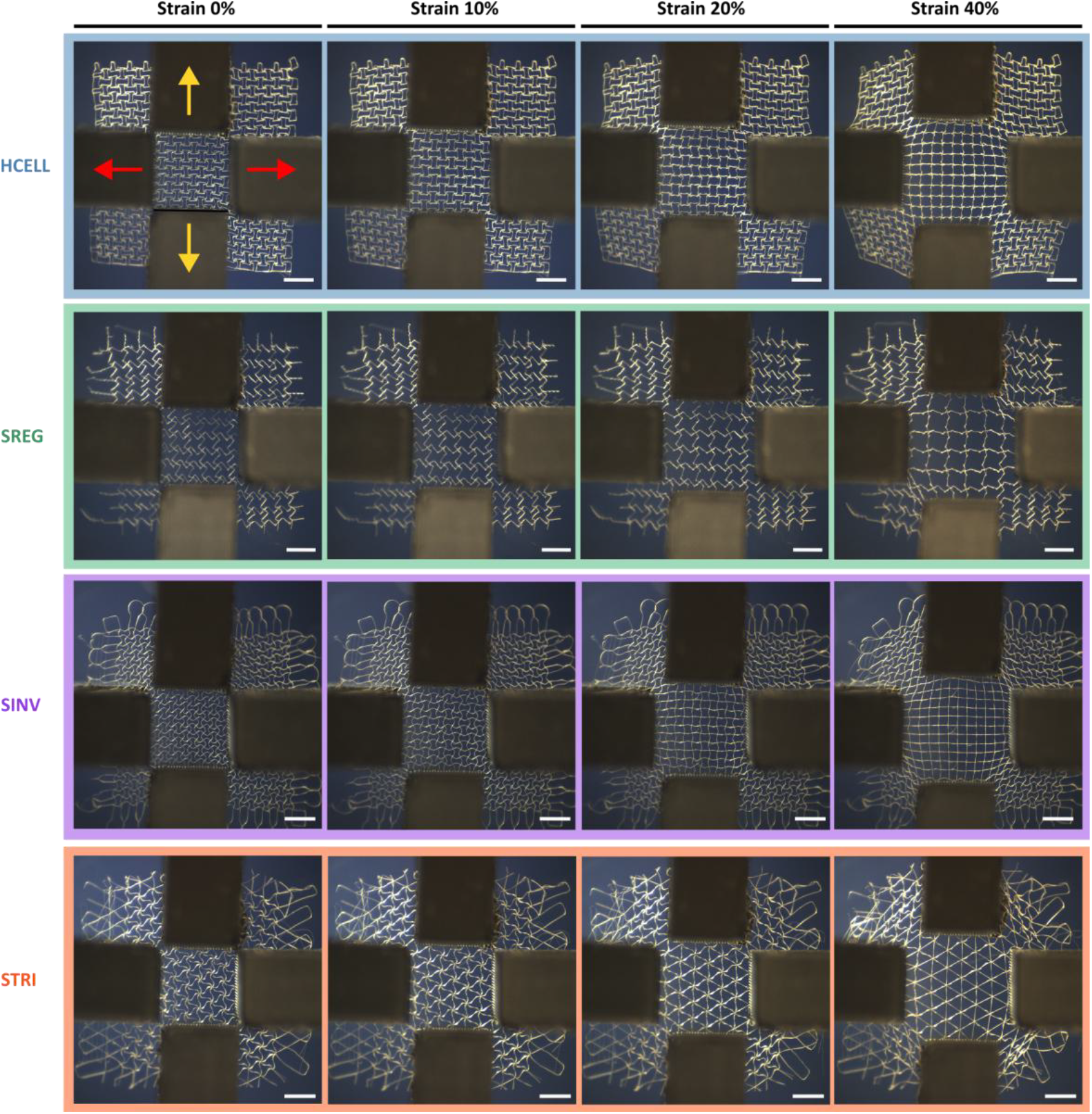
Panels showing scaffold deformation from 0% to 40% strain, exhibiting the transition from bi-directional unfolding to fully extended geometries. Red arrows show the direction of the tensile grips in the X axis, while yellow arrows represent the direction in the Y axis. (Scale bar = 2 mm)

### Computational model accurately matches auxetic scaffold mechanics

The numerical simulations were designed to replicate the auxetic behavior of the scaffolds during the biaxial tensile testing. The numerical prediction of geometry development aligned closely with the trends observed in the experimental tests. Additionally, the numerical simulation analysis of the *HCELL* scaffold provided displacement and stress maps, as shown in Figure 7. Based on the initial dimensions of the numerical model (5000 x 5000 microns), the strains of 0%, 20%, and 40% depicted in Figure 7 correspond to total displacements of 0, 500, and 1000 microns from the initial scaffold geometry configuration, respectively. The von Mises stress map highlights the stress experienced by the fibers during the simulation. At early stages, the stress values were low but progressively increased with higher strain levels, particularly in mechanically demanding areas such as the corners. In these regions, the fibers exhibited stress exceeding the defined PCL yield strength (12 MPa) in the numerical model, while the fibers within the scaffold interior showed minimal stress increases.

**Figure 7.**
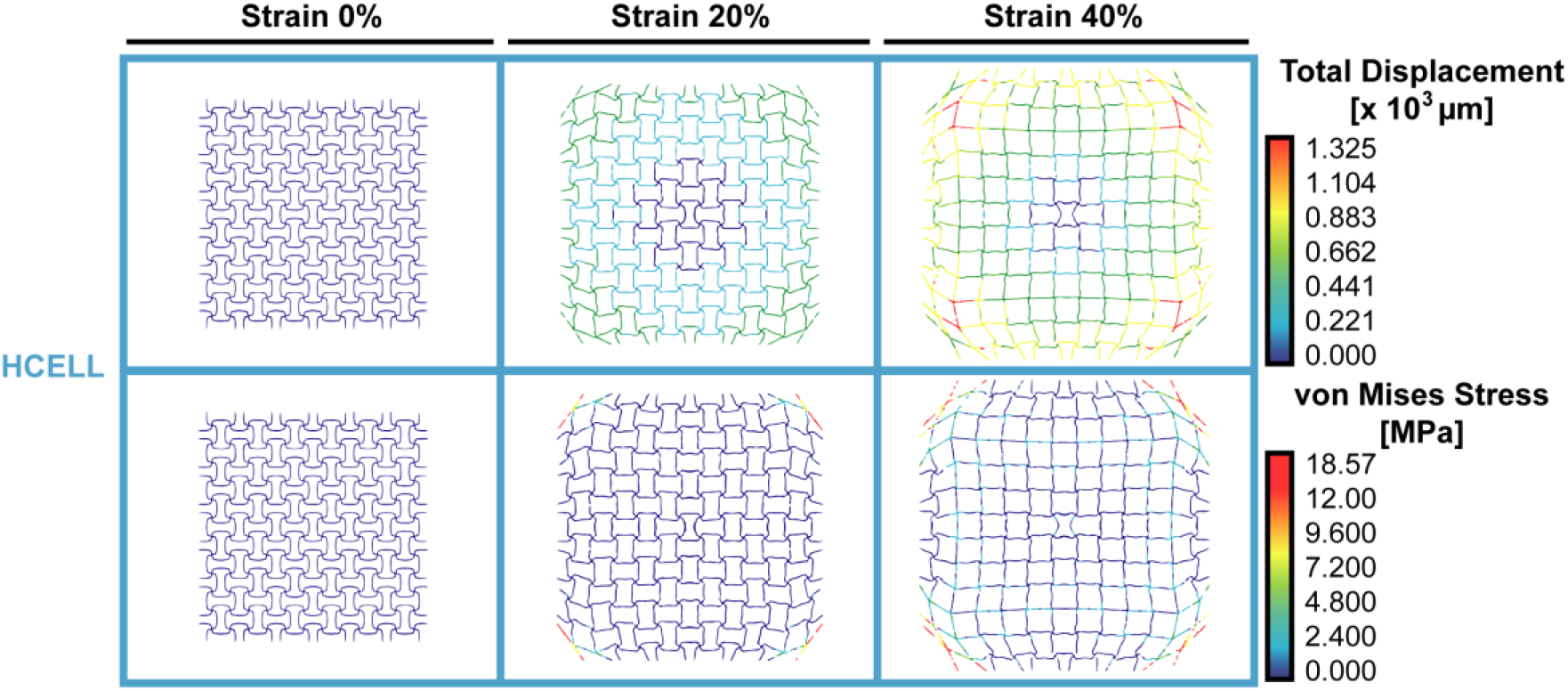
Sequential images from the HCELL numerical model capturing different deformation stages during the biaxial test simulation. Nodal total displacement and von Mises stress are illustrated within the scaffold geometry.

The non-linear response of the fiber scaffold is dictated by the auxetic microgeometry. Initially, the fibers unfolded with minimal resistance under mechanical load, allowing the scaffold to stretch easily. However, as deformation increased, the fibers gradually reached a fully extended state and tension started to rise exponentially. This transition marked what we previously denominated as the functional limit of the scaffold. For instance, in a hypothetical real-world application where the scaffold is attached to a wound, it would become significantly more challenging for the surrounding tissue to move in unison with the scaffold when subjected to strains beyond this functional limit. This mismatch could induce shear stresses, potentially causing scaffold detachment from the wound bed and interfering with biological processes necessary for proper wound healing, such as scaffold neovascularization. Therefore, careful consideration of this functional limit is important when designing auxetic scaffolds.

Hence, we simulated the biaxial tests up to deformations of 0.6, attempting to avoid excessively distorted geometry configurations that could lead to numerical artifacts (Figure 7). Figure 8 shows the stress-strain curves extracted from the numerical simulations of biaxial tensile tests and compared with their experimental counterparts.

**Figure 8.**
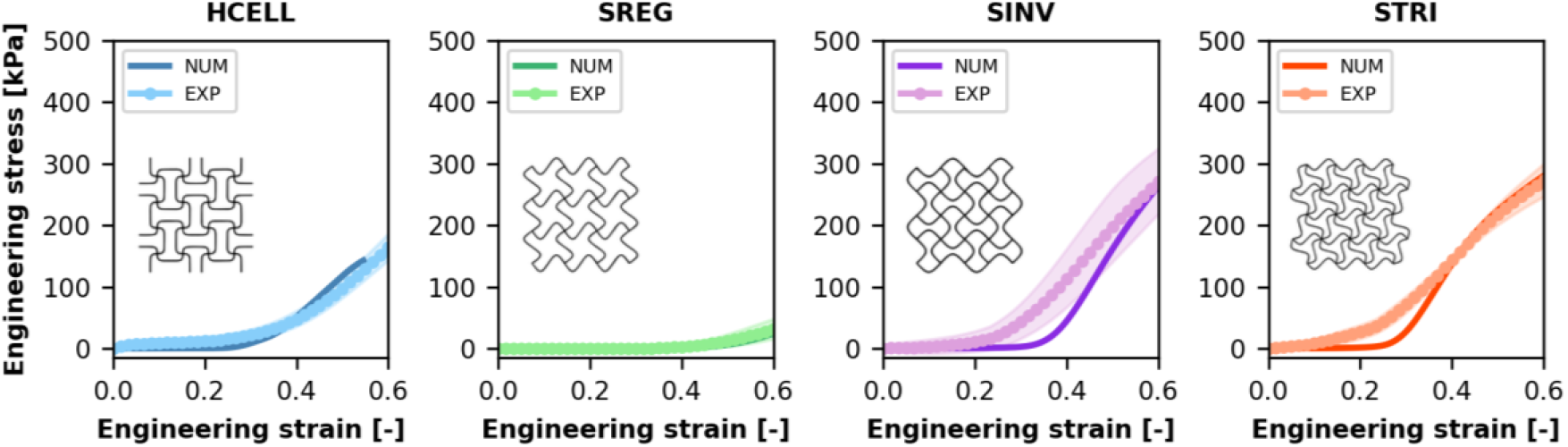
Engineering stress-strain curves of auxetic scaffold designs (HCELL, SREG, SINV, and STRI), comparing numerical simulation (NUM) and experimental test (EXP) results. Insets depict the auxetic design geometry of each scaffold. HCELL and SREG designs show strong agreement between numerical and experimental data, while SINV and STRI initially demonstrate a more compliant behavior in the numerical simulations that ends up converging with the experimental data at higher strains.

Computational models of *HCELL* and *SREG* scaffolds maintained soft auxetic behavior at strains up to 0.4 and 0.6, respectively (Figure 8), aligning excellently with the experimental data. These designs sustained a low stress response over a wide strain range, highlighting their capacity to withstand large deformations while exhibiting auxetic characteristics. In contrast, the simulations performed with *SINV* and *STRI* scaffolds reached earlier stages of strain, between 0.2 and 0.3, the stiffening associated with the non-auxetic behavior development. Numerical results from these simulations displayed a more compliant behavior at lower strain levels, resulting in lower stresses when compared to experimental data. However, at higher strains, numerical models began to reflect an increased stiffness, converging with the experimental test results.

The differences observed between the numerical and experimental results can potentially be attributed to variations in fiber intersections between the numerical model and the fabricated scaffolds. When fabricating these scaffolds via MEW, the molten material overlays and solidifies at the fiber intersections, forming thicker junctions with an increased cross-sectional area proportional to the number of intersecting fibers[51]. In the experimental tests, these junctions have demonstrated the ability to withstand considerable forces without failure, allowing deformations up to five times the original size of the scaffold. However, our computational model did not account for the thickened fiber intersections, which, we believe, are the origin of the observed discrepancies.

It is also important to note that the experimental tests exhibited considerable variability, which can be attributed to small yet cumulative geometric differences in each printed structure, even though they were derived from the same base G-code. To address this, we performed specific simulations that helped us correct the generated G-codes, as described in the following section.

### Altered mechanical behavior produced by MEW-induced distortions can be recapitulated and corrected via computational modeling

During the printing process, we identified geometrical distortions in several cases upon microscopic observation of the printed scaffolds, particularly in *SREG, SINV*, and *STRI* designs, where the printed structure did not match exactly the expected designed geometry. These discrepancies were caused by fiber lagging and layer shifting[52] when fiber deposition deviates from the printing trajectory forming the inconsistent vertical stacking of fiber walls, which has been reported previously[40]. To address this issue, we adjusted the printing path by slightly increasing the curvature trace for each layer in the G-code to compensate for the machine deviations[35]. This modification improved the fidelity of the fiber walls and allowed us to achieve geometries that closely matched the original auxetic design. From this point forward, scaffolds printed without this modification, thus presenting wall fiber deviations, will be referred to as “offset-printed scaffolds”, while the scaffolds with geometries matching the original auxetic design will be referred to as “normal-printed scaffolds”. It is important to note that, in the case of *SREG* scaffolds, instead of modifying the printing path, it was necessary to adjust the scale of geometric. Initially, we used a length of 125 µm for A and B parameters (Figure 2), but this resulted in low geometric fidelity and offset deviations. Thus, we decided to double these values to those shown in Table 1, which allowed us to obtain more precise structures without needing to alter the printing path.

Offset-printed scaffolds were also subjected to biaxial tensile tests to evaluate the extent to which these geometric discrepancies influenced the mechanical behavior of the scaffolds (Figure 9-A). As previously reported, the mechanical behavior in all cases begins with a soft response while the scaffolds retain their auxetic-designed geometry. However, with increasing strain, the fibers begin to unfold, causing the scaffolds to stiffen. Offset-printed scaffolds showed shortened upper layers during printing, causing the top fibers to fully extend at lower strain levels than the bottom fibers. Consequently, the stiffening of the structure also initiated at lower strains than in normal-printed scaffolds, a behavior consistent with the results observed in the biaxial tensile tests. This difference was especially pronounced in the *SREG* design due to the printing scale, as the *SREG* offset-printed scaffolds host more fibers within the structure for the same scaffold size. Thus, the different behavior in this case is caused by the number of fibers being involved rather than the geometry of the design.

**Figure 9.**
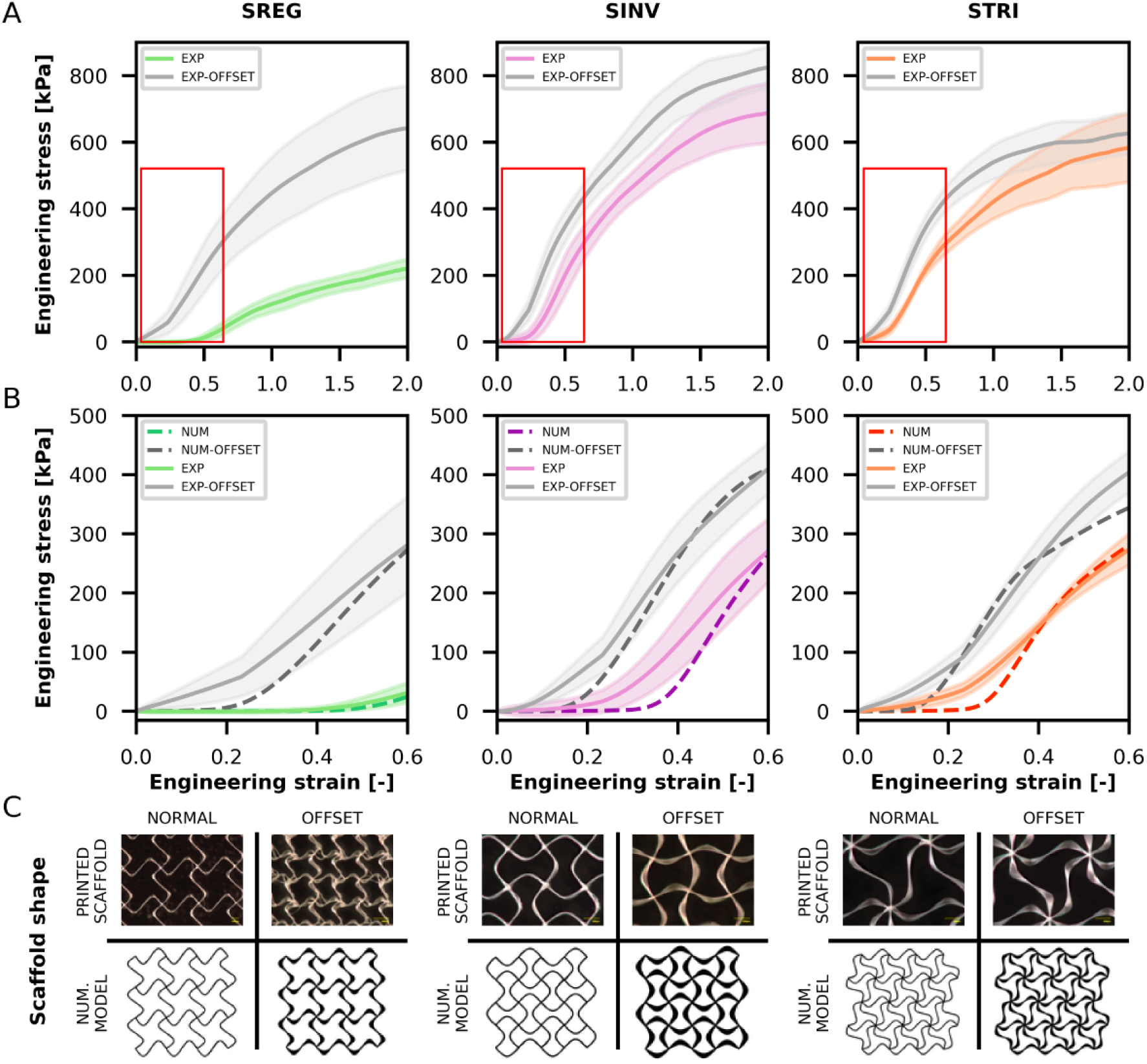
(A) Stress-strain curves extracted from the biaxial tensile tests of normal-printed and offset-printed scaffolds. Red boxes indicate the zoom areas of the plots located below. (B) Comparison between the results of numerical simulations and experimental tests of both normal-printed and offset-printed scaffolds within the auxetic strain range. (C) Details of normal and offset geometries observed in the printed scaffolds vs. normal and offset geometries designed in the numerical models.

Based on the observed geometry of the offset-printed scaffolds, we adapted our numerical model to better represent the offset-printed scaffolds with *SREG, SINV*, and *STRI* designs, aiming to capture the altered mechanical behavior exhibited by these scaffolds. Our analysis suggested that the shortened curvature observed in the offset-printed scaffolds could be simulated by decreasing parameter B (Figure 2). To implement this, we progressively reduced its initial value by 5% per layer in the code. While not perfectly, this modification successfully aligned the geometry of the numerical model with the experimentally observed, as shown in Figure 9-C.

While keeping all other parameters unchanged, we conducted numerical simulations using the new offset geometries and compared the results with the experimental tests performed on offset-printed scaffolds, as well as the numerical and experimental results from normal-printed scaffolds (Figure 9-B). Similar to the experimental findings, the stress-strain curves were translated, and the numerical models of the offset geometry scaffolds exhibited stiffer behavior compared to the normal geometry scaffolds.

Additionally, the comparison revealed a strong correlation between the numerical and experimental results for the offset geometries, showing the same minor discrepancies previously commented on and observed in the normal geometries (Figure 8).

### Auxetic MEW scaffolds exhibit mechanical behavior relevant to soft biological tissues

Given the biphasic, non-linear mechanical behavior exhibited by the fibrous auxetic scaffolds, we compared their performance with values reported for soft biological tissues. Although our primary goal was not to replicate the mechanical behavior of a specific tissue, skin serves as a useful reference due to the range of existing literature about its mechanical characterization. To evaluate scaffold biocompatibility, we performed normal human dermal fibroblast (NHDF) culture assays, which confirmed sustained NHDF viability over 7 days (Figure S5 – Supp. Info), indicating the scaffold’s suitability for soft tissue engineering applications..

It is important to note that the mechanical properties of soft tissues such as skin can vary widely depending on anatomical location, testing methodology, hydration state, and fiber orientation[53]. This variability poses challenges when attempting to define a single set of target mechanical values for scaffold design. Nonetheless, the auxetic scaffolds developed in this study fall within the lower-to-middle range of values reported for native soft tissues, suggesting their potential to accommodate various tissue-specific demands.

For example, Flynn et al.[54] reported that skin wound experience closure stress between 88 and 381 kPa during healing, which aligns with the stress levels supported by our scaffolds during deformation (Figure 9-B). In terms of stiffness, Annaidh et al.[55] found initial slopes of the stress-strain curve ranging from 0.7 to 1.5 MPa for back skin samples depending on orientation relative to Langer’s lines. Our scaffolds exhibit initial stiffness below 1 MPa (Figure 8), remaining in a comparable range. Furthermore, the failure strains of the scaffolds (Figure 5 and Figure 9-A) surpass 2.5, well within the broad range reported for human skin (0.16-2)[53], indicating substantial deformation capability without structural failure. In sum, with stress capacities in the range of 0–400 kPa, stiffness approaching 1 MPa, and strain-to-failure above 2.5, the auxetic scaffolds closely match or exceed the mechanical properties of native skin. In fact, their slightly reduced stiffness may be advantageous by minimizing mechanical mismatch at the tissue interface, thereby promoting better implant integration and comfort..

The auxetic fiber scaffolds developed in this study exhibit mechanical characteristics that not only align with values reported for skin but are also relevant to a range of soft tissues with similar nonlinear, anisotropic behavior. This highlights the potential of these designs as adaptable platforms in soft tissue engineering.

## 4. Conclusions

This study demonstrates the significant potential of auxetic PCL fiber scaffolds fabricated via MEW for soft tissue engineering applications. By combining mechanical testing and FEM simulations, we provide a comprehensive framework for designing scaffolds with tunable mechanical behavior.

All tested auxetic designs exhibited a characteristic biphasic mechanical response that closely mimics the non-linear behavior of soft tissues. Our computational model, built on a flexible beam framework, successfully predicted mechanical trends of both normal- and offset-printed scaffolds, validating its utility as predictive tool for scaffold optimization. This model also proved valuable in addressing fabrication challenges by enabling G-code corrections to mitigate fiber deposition offsets and ensure desired mechanical properties.

Importantly, the study revealed that even small deviations in printed design fidelity can impact scaffold mechanics, highlighting the role of manufacturing precision in MEW. *In vitro* cell culture experiments using normal human dermal fibroblasts (NHDFs) confirmed scaffold biocompatibility and supported cell adhesion and proliferation over a 7-day period (Figure S5 – Supp. Info), further supporting their suitability for tissue repair environments.

Nevertheless, our approach presents limitations. The computational model, while effective in predicting mechanical trends, does not fully account for complexities observed in the experimental tests such as the reinforced fiber intersections, which results in more compliant response at certain strain regimes. These differences are not very relevant in terms of stress magnitude but should be considered for real-world applications. Additionally, while we include plasticity effects, other material behaviors such as viscoelasticity or long-term degradation of PCL scaffolds, which may influence their performance *in vivo*, are not accounted for. Another limitation is that our testing scenarios focused solely on 2D stretching, which, while valuable, does not fully represent the complex, multidirectional forces scaffolds experience *in vivo*. Future work should include more comprehensive 3D mechanical testing and modeling to capture the behavior of auxetic scaffolds under physiologically relevant loading conditions, such as compression, shear, and combined loading. The biological assays were also limited to short-term evaluations using NHDF cells. Further investigations are necessary to assess the scaffolds’ long-term biocompatibility, degradation profiles, and integration within more complex *in vivo* models. Finally, while our study focused on PCL auxetic MEW scaffold designs, the applicability of our approach to other geometries, printing techniques, or materials remains unexplored. Expanding this work to include a broader range of designs and materials could provide a more versatile framework for future tissue engineering applications.

Looking ahead, several avenues for future research emerge from this study. For example, developing a regressive model could significantly streamline the design process by providing a predictive framework that directly correlates design parameters with mechanical and biological performance outcomes. Such a model could integrate machine learning or statistical regression techniques, trained on experimental and numerical data, to enable rapid, data-driven optimization of auxetic scaffolds. This approach would not only reduce the reliance on computationally intensive simulations but also allow for real-time adjustments during the design and fabrication phases. By incorporating manufacturing constraints, such as MEW process limitations or material properties, this tool could also predict feasible designs and generate optimized G-code directly, further bridging the gap between design and fabrication. Moreover, integrating biological data, such as cell proliferation or scaffold integration metrics into the model could provide a comprehensive optimization tool that considers both mechanical and biological performance. This holistic approach would be invaluable in developing next-generation scaffolds that are finely tuned for specific tissue engineering needs, ultimately accelerating the translation of auxetic scaffolds from research to clinical applications.

## Acknowledgements

Authors would like to acknowledge the Spanish Ministry of Economy and Competitiveness through the project PID2023-146072OB-I00. J.I. and M.C. acknowledge support from the Royal Thai Government for J.I.’s PhD scholarship (OCSC/MOST/MU/2562BE) and the Eindhoven Engine – BrainPort Eindhoven, through the AUXSTENT project. E.G.-G. is funded by a Ramon and Cajal Fellowship (RYC2021-033490-I, funded by MCIN/AEI/10.13039/501100011033 and the EU “NextGenerationEU/PRTR”). The authors declare that there is no conflict of interest.

