## Supplementary material for "Rational Design and Modeling of Auxetic Fiber Scaffolds for Soft Tissue Engineering via Melt Electrowriting": Supp. Info

**Supplementary Information**

**Melt electro-writing set up**

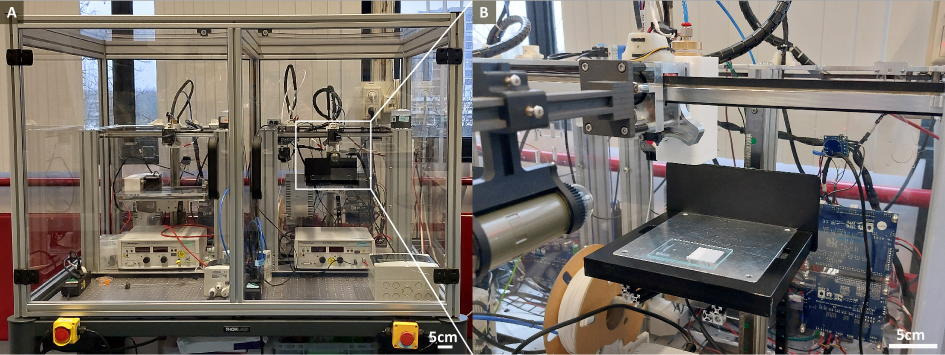

Figure S1: The machine set up for melt electro-writing. Motion system and enclosure were installed on the optical table (A). Close-up image showing the Dinolite camera, printing cartridge, moving head (X&Y axis), collector (Z axis), and specimen on the glass slide (B).

**Uniaxial tensile test for PCL characterization**

The specimens were made by melting medical grade PCL (PC12, Corbion, Netherlands) into the casting mold with dog bone shape (ASTM D638 type V), made from Polydimethylsiloxane (PDMS) (Figure S2-A). The gauge length ($L_{0}$) of the specimens was assumed to be 9.73mm, while the width ($W$) and thickness ($T$) of the specimens were measured before the test. The bulk properties of PCL were characterized by uniaxial tensile machine (MTS criterion 42) with five specimens (Figure S2-B, C). The tensile stress ($\sigma$) and strain (ε) were calculated from equation 1 (Eq. 1) with the force ($F$) recorded from 500N load cells, and the displacement ($L$) from the tensile machine with a strain rate of 10 mm/min. The yield point was defined by a maximum ε in the linear part of elastic region before plastic deformation identified by least-square regression (R^2^>0.99). The stiffness ($E$) was then calculated based on regression’s coefficients from this linear part and strain energy density was derived from the area under $\sigma-\epsilon$ curve to the yield point, as shown in Figure S2-D.

$$\sigma=F/ WT$$

(1)

$$\varepsilon=\frac{L}{L_{0}} -1$$

(2)

The width of specimens (5 replica) before and after tensile tests were 3.78 ± 0.13, and 1.54 ± 0.12 mm, respectively. After the test at an engineering strain of 534%, all specimens plastically deformed without breaking or rupturing. The calculated Poisson’s ratio is 0.10 ± 0.01. Table S1 presents the average (± standard deviation) of stiffness, strain energy density (U_s_), strain and stress at yield point, and ultimate tensile strength (UTS) that calculated from tensile result (Figure S2-E).

| **Stiffness [MPa]** | **U_S_ [MJ/m^3^]** | **Yield strain [%]** | **Yield stress [MPa]** | **Strain at UTS [%]** | **UTS [MPa]** |
| --- | --- | --- | --- | --- | --- |
| 97.23 ± 7.57 | 0.71 ± 0.18 | 0.12 ± 0.02 | 10.89 ± 1.54 | 0.23 ± 0.02 | 14.03 ± 1.48 |

Table S1. Bulk properties of PCL extracted from uniaxial tensile test and implemented on computational model.

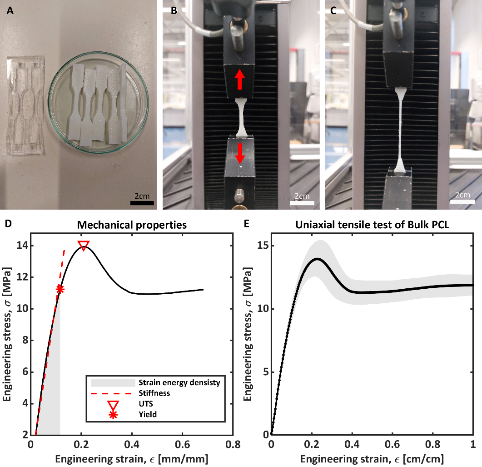

Figure S2: Uniaxial tensile test and mechanical properties of bulk PCL. Casting mold with dog bond shape and specimens (A). The specimen before (B) and after (C) the tensile test, where the red arrows represent the direction of tensile strain. The quantification of yield point, ultimate tensile strength (UTS), stiffness, and strain energy density (D). The σ-ε curve of bulk PCL specimens where a black line shows an average of five specimens and a grey shade represents a standard deviation (E).

**Python-based application for G-code generation**

The scaffold geometries were designed in Python (Python 3.12), allowing users to customize the geometry through parameter A and B. To correct the layer shifting and offsetting during fabrication[1], the program incorporates phase and amplitude adjustments reported in previous work[2] for effective correction. By providing geometry customization and addressing fabrication errors, this program enhances control and reproducibility, thereby facilitating the extension and exploration of new scaffold designs. This program relies on standard Python libraries and operates without the need for additional commercial software or runtime, making it accessible and cost-effective for researchers and practitioners. In addition, the program produces G-code (Duet3D, RepRapFirmware 3.4) that is widely used in 3D printers and especially in the open source MEW platform[3]. This direct G-code generation streamlines the workflow by eliminating the need for intermediate software to translate the designed geometry into machine instructions.

**G-code Application workflow**

Step 1: Install Python (version 3.12 or later)

Step 2: Choose the python script from the scaffold design in Gcode folder

- HCELL design: Gcode/S-regular/generate_h_cell.py. (P05 design [1])
- STRI design: Gcode/S-regular/generate_s-triangular.py
- SINV: Gcode/S-regular/generate_s-inverted-small.py (P02 design [1])
- SREG: Gcodes/S-regular/generate_s-regular-large.py (P03 design [1])
- Arrowhead: Gcode/Arrowhead/generate_arrowhead.py

Step 3: Modify variables in the python script

| Design | Variable | Description |
| --- | --- | --- |
| HCELL | A | Unit cell variable (mm). See the original manuscript for more detail |
|  | B | Unit cell variable (mm). See the original manuscript for more detail |
|  | alpha | Scaling factor for B. Used to change the aspect ratio of the unit cell |
|  | X_repetitions | The number of unit cells in X axis |
|  | Y_repetitions | The number of unit cells in Y axis |
|  | CTS | Critical translational speed (mm min^-1^) |
|  | layers | The number of layers |
|  | number_scaffold | If >1 the script will generate a G-code file for printing >1 scaffold at the same time. This will increase the total size of the printing area. Please check that the total size matches the size of glass slide or the collector. |
|  | offset_bw_scaffold | Offset (mm) between scaffolds. Used when number_scaffold > 1 |
| STRI | A,B, … | All variables in HCELL except alpha |
|  | zero_degrees_length | Extra distance (mm) before U-turn to prevent unstable jet due to a sharp u-turn. |
|  | sixty_dg_length | Same as above but for ± 60° lines |
| SINV | A,B, … | All variables in HCELL except alpha |
|  | uturn_length | Extra distance (mm) before U-turn to prevent unstable jet due to a sharp u-turn. |
|  | uturn_radius | Radius of U-turn (mm) |
| SREG | A,B, … | All variables in SINV |
| Arrowhead | A,B, … | All variables in HCELL |
|  |  | The toolpath offset, layer shifting, and printing speed correction described in [1] can be modified in Gcode/Strategy/layer_correction.py |

Step 4: Run python script

Step 5: (Optional) Visualize the generated G-code file

- Go to <https://ncviewer.com/> and upload the G-code file
- Alternatively, we developed a custom web application to visualize G-code at <https://gcode-visualizer.pages.dev/> and released the source code in Github (<https://github.com/nodtem66/gcode_visualizer>)

**Abaqus FEM model scripts**

The numerical model was developed with Abaqus software v6.14 (3DS Dassault Systems). The main script to create the FEM model and start running simulation is located at FEM/gen_inp.py. The following command line shows an example of the script usage. The details of geometry, material property, and boundary conditions are described in the script FEM/create_aux_str.py.

| > | abaqus cae noGUI=gen_inp.py -- design a b d xr yr zr id |
| --- | --- |

| Parameter | Type | Description |
| --- | --- | --- |
| design | String | The value must be one of the following keywords:   - arrowhead - hcell_uniform - stri_uniform - sreg_uniform - sinv_uniform - stri_shift - sreg_shift - sinv_shift   The second part of the keyword after underscore is the variation of geometry where uniform means that the fiber walls stack vertically and shift means the fiber walls stack with a shift or an offset between layers. |
| a | Float > 0 | Geometry parameter A. It has a different meaning for each design, but it is always defined in microns (10^-3^ mm). |
| b | Float > 0 | Geometry parameter B, defined in microns except for the arrowhead design where it is defined as an angle (°) |
| d | Float > 0 | Fiber diameter in microns (10^-3^ mm). |
| xr | Integer | Number of repetitions of the basic design unit in the x-direction |
| yr | Integer | Number of repetitions of the basic design unit in the y-direction |
| zr | Integer | Number of layers stacked in the z-direction |
| id | Integer | Unique numerical identifier, starting from 00001 |

**Arrowhead design, fabrication, and characterization**

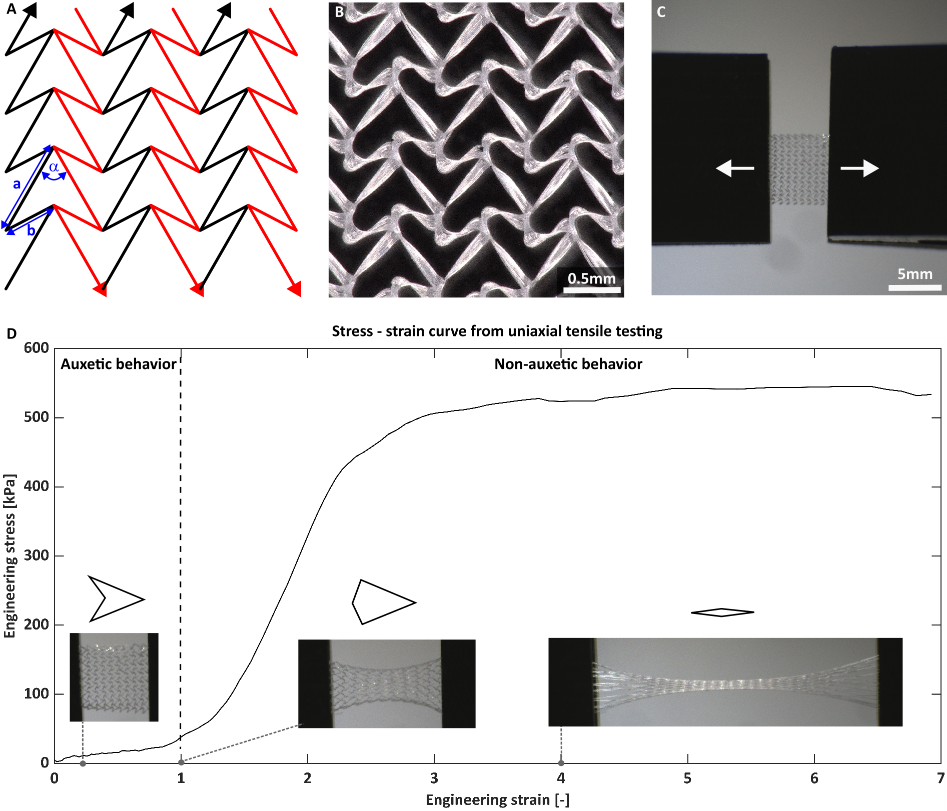

Figure S3: Fabrication of arrowhead scaffold and uniaxial tensile testing. Printing schematic diagram, showing the printing path as black and red arrows where a = 1 mm, b = 0.5mm, and α = 45° (A). The image of fabricated scaffold from digital microscope (B). The scaffold before uniaxial tensile testing where the white arrows show the direction of tensile force (C). The arrowhead scaffold exhibited auxetic behavior in 0 – 1 strain before turning to non-auxetic behavior (D).

**Scaffold pore size and area calculations**

Figure S4: The geometry of pores derived from G-code position for calculating theoretical pore size and pore area. Prior to the calculation, the geometry (solid color) was offset inward to remove the area occupied by surrounding fibers (dash lines). The theoretical pore areas of HCELL (blue), SREG (yellow), SINV (green), STRI (red) are 0.257, 0.920, 0.215, and 0.383 mm^2^, respectively while the theoretical pore sizes (minimal ferret diameter) of these designs are 0.523, 1.226, 0.460, and 0.917 mm, respectively.

**Cellular viability evaluation**

**Materials and Methods**

**Cell culture on scaffolds**

Due to the hydrophobic nature of PCL, a surface hydrophilization treatment had to be performed the day before cell seeding. In this process, samples were immersed in a 5M NaOH solution at room temperature for 5 hours.[4,5] Subsequently, the solution was removed, and the samples were washed with phosphate buffered saline (PBS, Thermo Fisher Scientific, MA, US). Finally, the samples were sterilized by immersion in absolute ethanol and exposed to UV light for 20 minutes on each side.

Normal human dermal fibroblasts (NHDF, Lonza, Switzerland) were cultured under standard conditions (5% CO_2_, 37ºC) up to 70-80% confluence in their regular expansion medium: Dulbecco’s Modified Eagle Medium (DMEM) high glucose (4.5 g/L) supplemented with: 10% fetal bovine serum (FBS, Thermo Fisher Scientific, MA, US), 100 U/ml penicillin, 100 µg/ml streptomycin, and 2 mM L-glutamine (all from Lonza, Switzerland). For cell expansion, cultures were washed with PBS, detached with TrypLE^TM^ Express (Thermo Fisher Scientific, MA, US) and plated in T25 cell culture flasks at a density of 15,000 cells/cm^2^.

A cell-laden collagen hydrogel was prepared with a final collagen concentration of 2.5 mg/mL. The process involved diluting a Rat Tail collagen type I solution (stock 10.8 mg/mL, Corning, NY, US) in an ice bath, incorporating the previously prepared cell suspension, 10x Dulbecco’s phosphate-buffered saline (DPBS), and 0.5 M NaOH (both from Sigma-Aldrich, Germany) to adjust the pH to 7.4. The NHDF cells were ultimately suspended at a concentration of 10^6^ cells/mL in the collagen solution. Subsequently, 20 µL of the solution was gently pipetted onto the scaffold piece (4x4 mm) and placed in a humid chamber at 37ºC for 20 minutes to crosslink and solidify the collagen solution. Upon gelation, the hydrogel was hydrated with DMEM, with the medium being changed every 48 hours to ensure continuous nutrient delivery to the cultures.

**Fibroblasts viability**

The Alamar Blue assay was employed to assess cell viability over a 7-day culture period. Alamar Blue reagent (Thermo Fisher Scientific, MA, US) was diluted in the culture medium and added to the sample wells (Alamar Blue 1:10 DMEM). The solution was then incubated for 4 hours under standard culture conditions (37°C and 5% CO_2_). Subsequently, samples of the solution were collected, and fluorescence was measured at excitation 560 nm/emission 590 nm using a spectrophotometer (Synergy HT Multi-mode microplate reader, BioTek Instruments, VT, US). Cell viability was monitored starting from day 1 of the culture and subsequently at 48-hour intervals. Cell-laden collagen hydrogel incubated for 7 days in well plates was used as a control. Every condition was conducted in triplicates.

**SEM inspection**

Scanning electron microscopy (SEM) was conducted on the samples to corroborate collagen laden within the scaffold. Moreover, collagen-based hydrogel morphology around the PCL fibers was observed.

After the fixation steps, the samples were subjected to sequential dehydration in graded series of ethanol. Subsequently, they carried out a total water adsorption procedure by submitting them to a critical point drying stage (Leica EM CPD300 Critical Point Dryer, Germany). Finally, the samples were coated with a 20 nm carbon film before they were examined by SEM. Then, samples were visualized at 10 kV and 3 spot sizes, using a field emission SEM Inspect^TM^ F50 (FEI, OR, US) at 10 keV.

For the acquisition of the collagen layer thickness electron images, a Dual-Beam Nova Nanolab 200 (Thermo Fisher, MA, US) was used. This equipment features a Ga+ ion beam, which facilitates the growth and milling of materials, thereby enabling the observation and acquisition of secondary electron images from the material’s interior. Initially, a platinum (Pt) layer was deposited via electron beam-induced deposition (EBID) to protect the surface of the area to be analyzed from ion irradiation damage. Subsequently, an additional Pt layer was deposited using ion beam-induced deposition (IBID). Following the Pt deposition, coarse milling was performed using a regular cross-section with a beam current of 0.3 nA. Subsequent cleaning steps were conducted by progressively reducing the beam current down to 50 pA.

**Cell morphology visualization**

Morphological analysis of NHDF cells was performed at the end of the cultures. Cells were fixed inside the sample wells culture using 4% (w/v) paraformaldehyde (PFA, Thermo Fisher Scientific, MA, US) for 30 minutes, blocked overnight in PBA with 5% bovine serum albumin (VWR, PA, US) and incubated overnight with Phalloidin-TRITC (0.1 mg/mL in PBS) and DAPI (0.01 mg/mL in PBS), both from Sigma-Aldrich (Germany). Samples were observed with a confocal microscope (Zeiss AXIO observer, Zeiss, Germany). Maximum intensity and 3D volumetric images were generated from z-stacks using ImageJ software (MD, US).

**Statistical analysis**

For fibroblast viability, after data normality assessment via quantile-quantile and normal probability plots, analysis of variance (ANOVA) followed by post hoc Tukey-Kramer tests was performed to determine statistical significance among the studied continuous variables in the different conditions. A non-parametric (Kruskal-Wallis) test followed by post-hoc Tukey-Kramer tests was used instead when data distribution was not normal.

**Results**

**NHDF cells are viable and proliferate over seven days in the fiber scaffold**

To assess the in vitro biocompatibility of the scaffold and its ability to support cell growth, we cultured NHDF cells within a collagen-based hydrogel homogeneously and well distributed across the surface of the fibers of *Arrowhead* designed scaffolds (Figure S5-A, B). Embedding the cells in a collagen I hydrogel ensured that the cells were evenly dispersed throughout the biomaterial during the entire cultivation period. Importantly, the cells adhered to the scaffold fibers and did not remain solely within the hydrogel filling the pores of the auxetic structure (Figure S5-C).

The viability was measured using the Alamar Blue metabolic assay (Figure S5-D). Results showed an initial increase in metabolic activity during the first few days, suggesting that the cells adapted well to their environment and proliferated effectively. This early phase of increased activity indicated healthy cell growth and successful attachment to the scaffold. The stabilization of metabolic activity by day 7 indicated that the cells had reached a steady state, maintaining their functions without signs of stress or decline. This highlights the scaffold’s ability to provide a stable environment for cell growth, a key factor for its potential use in a clinical application.

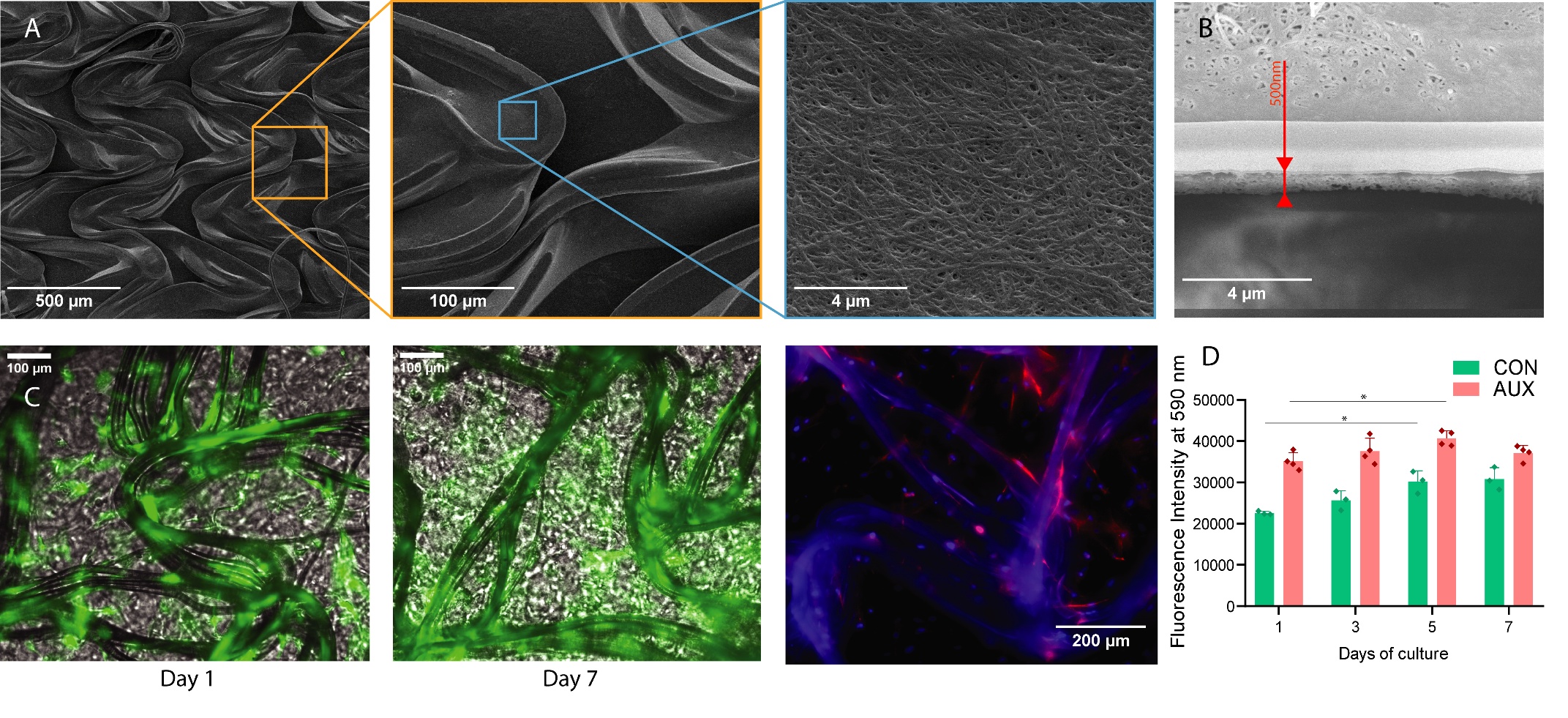

Figure S5. (A) Representative SEM images of a PCL auxetic fiber scaffold seeded with collagen-based hydrogel. (B) Transversal cut of the collagen layer by Dual beam SEM and tilted 52º. (C) GFP-NHDF (Green) in the auxetic scaffold after 1 and 7 days of culture. Dapi-phalloidin staining of the auxetic fiber scaffold seeded with NHDF after 7 days of culture. Cell nuclei appear in blue (dapi), while cell cytoskeleton morphologies appear in red (phalloidin). (D) NHDF viability over time within collagen hydrogel culture in well plates (CON) and within the PCL auxetic scaffolds (AUX). Triplicates were analyzed and p<0.05 was considered for statistical significance.
